# Aggregation-Prone Region Mapping in Olfactomedin Domain of Myocilin through Classical and Enhanced Sampling Molecular Dynamics Simulations

**DOI:** 10.1101/2025.08.02.668245

**Authors:** Inci Sardag, Zeynep Sevval Duvenci, Emel Timucin

## Abstract

The aggregation of the myocilin olfactomedin (OLF) domain, generally driven by genetic mutations, is the leading cause of primary open-angle glaucoma (POAG). Developing therapeutic strategies requires a detailed understanding its initial unfolding events that expose aggregation-prone regions (APRs). However, it has been a challenge, as the slow conformational dynamics of OLF hinders classical molecular dynamics (MD) simulations from capturing aggregation-prone OLF intermediates. To overcome this, we employed a multi-pronged computational strategy, integrating over 15 *µ*s of simulation time across diverse conditions, including high-temperature, enhanced sampling, chemical denaturation, and simulations of the pathogenic I499F mutant. Our results reveal that OLF unfolding is not random but initiates at specific structural regions pertinent to the terminal blade A and E. Specifically, the blade interfaces between A-B and A-E showed unique regions rich in aromatic/hydrophobic residues as aggregation hotspots. Overall, our simulations proved effective to generate a detailed map of seven distinct APRs. The accuracy of these APRs is partially validated by the close localization of these predicted regions with both previously identified amyloid peptides and the sites of known disease-causing mutations. By scrutinizing the OLF structure and dynamics under different MD settings, our study provides potential molecular targets for developing new therapeutic interventions against POAG.

## Introduction

Proteins fold into three-dimensional structures to perform their functions correctly. Mutations and/or environmental stressors can disrupt this folding, leading to misfolding and aggregation [1]. One form of aggregation is amyloid formation, wherein misfolded proteins stack to form highly ordered fibrillar structures [2]. This occurs when typically buried segments of proteins, known as amyloid-prone regions (APRs), become exposed, facilitating the intermolecular interactions that produce the characteristic fibrils [3]. Amyloid formation is a hallmark of numerous diseases, spanning neurodegenerative and systemic disorders, making it a central focus in both basic and applied biomedical research [4–8].

Among the catalog of amyloid-forming proteins, myocilin has emerged as a relatively novel example [9, 10]. Myocilin comprises an N-terminal coiled coil domain containing a leucine zipper motif, a C-terminal olfactomedin domain (OLF), and an unstructured linker region connecting these termini [11, 12]. Myocilin undergoes proteolytic cleavage within the linker, releasing OLF into the extracellular matrix (ECM) [13–15]. Mutations in the OLF domain are associated with primary open-angle glaucoma (POAG) [12], wherein a toxic gain-of-function mechanism is widely accepted, proposing that mutant myocilin forms insoluble aggregates that impair secretion and elevate intraocular pressure [16–18]. While a loss-of-function mechanism is considered unlikely, given that individuals with myocilin truncations do not develop POAG [19, 20]. Pathogenic mutations that lower thermal stability of OLF promote amyloid formation at physiological conditions [9, 21]. Notably, even wild-type (WT) OLF can form amyloid aggregates when exposed to mildly destabilizing conditions, adopting non-native conformations that readily fibrillize at elevated temperatures [9, 21].

The OLF domain is a 30 kDa protein with a five-bladed *β*-propeller fold, in which the blades radiate symmetrically around a central axis (Fig.1) [11, 22, 23]. It adopts a toroid structure formed by the interactions among the five inter-blade surfaces. Each blade is composed of four antiparallel *β*-strands extending outward, connected by loops and/or surface helices. Terminal *β*-strands fold over one another, creating a molecular clasp that stabilizes the toroid (Fig.1). C245 in blade E forms a disulfide bond with C433 in blade D [11, 24]. OLF domain houses a hydrophilic dimetallic center that binds Ca^2+^ and Na^+^ ions [25]. A long loop between residues E359 and I379 forms a valve-like structure at the BC interface. Ca^2+^ is coordinated by D380, N428, A429, I477, and D478, while Na^+^ localization involves G326, L381, and D478. Together, the five-bladed *β*-propeller fold, the molecular clasp at the blade termini, the C245–C433 disulfide bond, the hydrophilic dimetallic center, and the long loop spanning blades B and C define critical structural features of the OLF domain (Fig. 1).

**Fig. 1.**
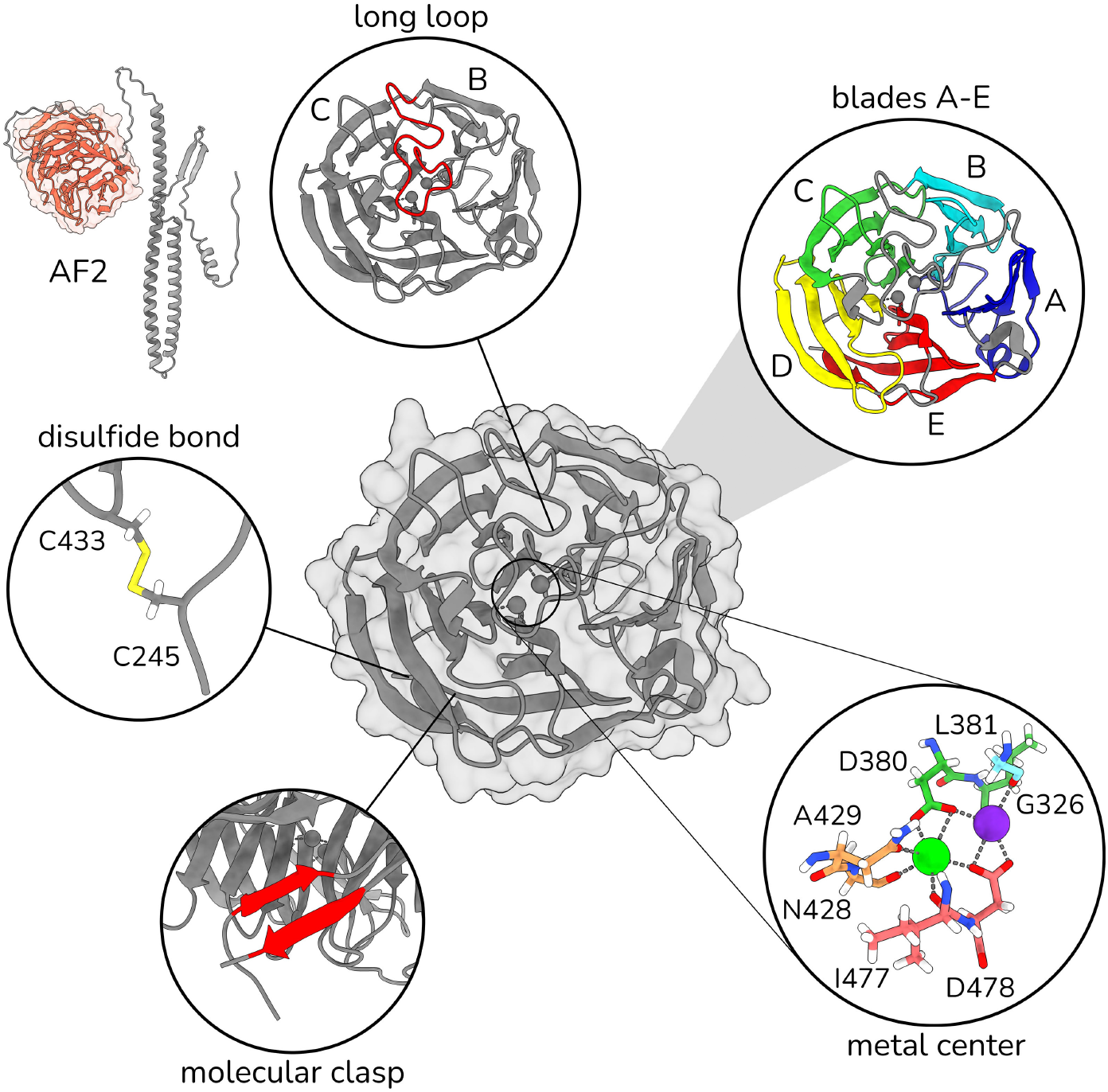
Important structural motifs of the OLF domain of human myocilin (PDB id: 8frr).

Previous molecular dynamics (MD) simulations of OLF showed that its structural fluctuations are largely restricted to loop regions and the metal center over a 10-*µ*s timescale [10]. Since APR exposure often results from unfolding or large domain motions, the limited dynamics captured at the microsecond scale suggest that longer simulations or enhanced sampling methods are needed to fully map its conformational landscape and amyloidogenic potential. In this study, we employ multiple MD methodologies to explore how the OLF domain transitions from a folded to a partially unfolded, aggregation-prone conformation. Our aim is to uncover aggregation-prone regions (APRs) and characterize intermediate states of OLF linked to primary open-angle glaucoma (POAG). To achieve this, we utilized Gaussian-accelerated molecular dynamics (GaMD) with varying boost energies, alongside classical molecular dynamics (cMD), to enhance sampling of OLF conformations. Additionally, we investigated various denaturant conditions such as elevated temperatures, urea concentrations along with the severe POAG mutant of I499F using cMD. Through the integration of all-atom MD simulations under diverse settings, this work provides insights into the structural mechanisms that underlie possible OLF conformations driving amyloid formation.

## Methods

### OLF Structure

All structures of the OLF domain of myocilin in PDB were retrieved from PDB as of January 2025. A total of 26 PDB structures of OLF domain were analyzed with principal component analysis (PCA) based on Cartesian coordinates of C*α* atoms using Bio3D [26]. The recent WT OLF structure with the PDB id 8frr [10] was selected as a representative of the largest cluster with fifteen members (Fig. S1).

### MD Simulations

Buffer contaminants and water molecules were removed from the the 8frr structure, while two metal ions (Ca^2+^ and Na^+^) were retained within the hydrophilic central cavity (Fig. 1). Protonation was done using a neutral pH of 7.0 by PDB2PQR [27]. The protonated OLF structures was placed in the center of orthorhombic boxes with an edge distance of 10 Å from the protein. Solvated systems were neutralized by adding Na^+^ and Cl^*−*^ at a final concentration of 0.15*M*. All MD systems were prepared using CHARMM-GUI [28–30], except for the urea-containing systems, which were prepared by PACKMOL [31, 32]. Two urea solvated systems were generated using the 1200:11,000 and 2400:10,000 urea:water ratios. Resulting MD systems were energy minimized for 10,000 steps followed by a 250 ps NVT equilibration step at 298K. An NPT ensemble was used for all production simulations. Pressure was kept at 1 atm and temperature at 310, 365 or 410 K using the Langevin pressure and temperature coupling [33– 35]. NAMD (v3) algorithm was used for all MD simulations [36] with the CHARMM36m force field [29, 37, 38]. Water molecules were explicitly treated with the TIP3P model [39]. For all simulations, periodic boundary conditions and an integration time step of 2 fs were used. Long-range Coulomb interactions were calculated using the Particle-Mesh Ewald (PME) method with 1 Å grid spacing [40]. A cut-off of 12 Å was used for non-bonded interactions. A total of 15 OLF simulations were performed, differing in simulation temperature, solvent type, mutation status, presence or absence of the C245-C433 disulfide bond, and boost potential (Table 1). Details of the resulting MD systems is summarized in Table S1.

**Table 1.** MD System and Simulation Parameters.

| OLF state | method | Disulfide bond | solvent | effect |
| --- | --- | --- | --- | --- |
| WT | cMD (310K) | reduced | water |  |
| WT | cMD (365K) | reduced | water | thermal stress |
| WT | cMD (410K) | reduced | water |  |
| WT | cMD (365K) | oxidized | water |  |
| WT | cMD (410K) | oxidized | water |  |
| I499F | cMD (310K) | reduced | water | thermal stress |
| I499F | cMD (365K) | reduced | water | incread KE |
| I499F | cMD (365K) | oxidized | water | POAG variant |
| WT | cMD (310K) | oxidized | urea | chemical<br>denaturant |
| WT | cMD (310K) | oxidized | urea |  |
| WT | cMD (310K) | reduced | urea |  |
| WT | GaMD (dihedral) | reduced | water | boost<br>potential |
| WT | GaMD (dual) | reduced | water |  |
| WT | GaMD (total) | reduced | water |  |
| WT | GaMD (total) | oxidized | water |  |
| cMD: classical MD, GaMD: Gaussian accelerated MD, KE: kinetic energy |  |  |  |  |

### Gaussian Accelerated MD Simulations

Enhanced sampling was performed using Gaussian accelerated molecular dynamics (GaMD) [41, 42]. A harmonic boost potential (Δ*V*), following a Gaussian distribution, is introduced into the system when the original potential (*V* (*r*)) is below a threshold energy (*E*):

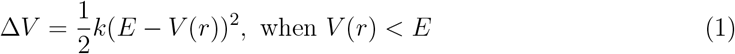

where *k* is the harmonic force constant. The modified potential *V* ^*∗*^(*r*) is then given as:

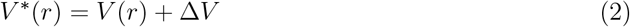

When the original potential is above the threshold energy (*E*) (*V* (*r*) *> E*), then the boost potential (Δ*V*) is set to zero, leading to *V* ^*∗*^(*r*) = *V* (*r*). GaMD was applied in three distinct modes: (i) total boost mode, in which the threshold energy (*E*) is set based on the average total potential energy; (ii) dihedral boost mode, in which *E* is set based on the average dihedral potential energy; (iii) dual boost mode, in which both total and dihedral boost potentials are applied simultaneously. Before GaMD production runs, an additional equilibration was performed for 1 ns to collect threshold energies and other parameters.

### Trajectory Analysis

Production trajectories were analyzed by pairwise root mean square deviation (RMSD) and fluctuation (RMSF) of C*α* atoms using MD analysis and in-house scripts [43, 44]. Solvent-accessible surface area (SASA) changes of the OLF domain and five A-E blades were calculated. Radius of gyration (*R*_*G*_), radial distribution function and atom-atom distances were computed using VMD. For the visualization of structures and trajectories, VMD and ChimeraX were used [45–48]. Fraction of native contacts (Q) was calculated with a soft cutoff that uses a sigmoid function [49]:

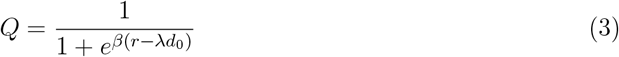

wherein; *d*_0_ is the distance in the native state, *β* controlling the steepness of the transition is set to 5.0, and *λ* defining the tolerance for deviations from the native distance is set to 1.8. The MDAnalysis function was used in this analysis [43, 44].

### Free Energy Landscape Analysis

MD trajectories were analyzed using PCA based on the cartesian coordinates of C*α* atoms. The covariance matrix *C*_*ij*_ was computed as follows:

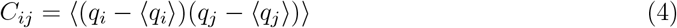

where *q*_*i*_ and *q*_*j*_ represent the Cartesian coordinates of atoms *i* and *j*, and ⟨…⟩ denotes the ensemble average over the entire trajectory. The eigenvectors and eigenvalues were obtained through diagonalization of the covariance matrix, with the eigenvectors representing the principal components (PCs) and the eigenvalues indicating the magnitude of fluctuations along these components.

The free energy landscape (FEL) analysis was carried out by projecting the MD trajectory onto the first two PCs. The probability of finding the system in a particular state *q*_*x*_ of the reaction coordinates (in this case, the principal components) is defined proportional to the free energy of the state as follows:

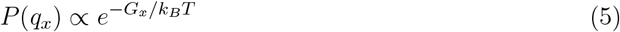

where *G*_*x*_ is the free energy of the state *q*_*x*_, *k*_*B*_ is the Boltzmann constant, and *T* is the temperature. The free energy landscape is then obtained by the given relation:

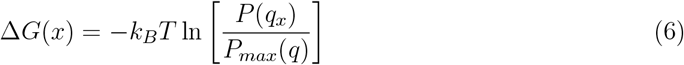

where *P* (*q*_*x*_) is the probability of finding the system in the state *q*_*x*_ in the PC1-PC2 space, *P*_*max*_ is the probability of the most populated state.

### Structure-based Aggregation Propensity Prediction

Based on FEL analyis, the minimum energy state of the MD trajectories were selected from each system. For trajectories showing more than one minima on FEL, a distinct structure is selected for each minimum. The crystal structure and the selected OLF conformations from MD trajectories were analyzed using aggregation propensity prediction tools, namely AggreScan3D [50–54] and CamSol [55, 56]. Adaptive Poisson-Boltzmann Solver (APBS) was used to analyze surface electrostatic potential distributions of selected MD structures [27].

## Results

### Assessing MD-induced Structural Changes in the OLF and its Individual Blades

To evaluate the extent of conformational changes in the overall OLF structure and its individual blades, we determined the fraction of native contacts (Q) (Fig. 2a), employing a soft cutoff for its ability to more accurately represent the continuous nature of interactions [49].

**Fig. 2.**
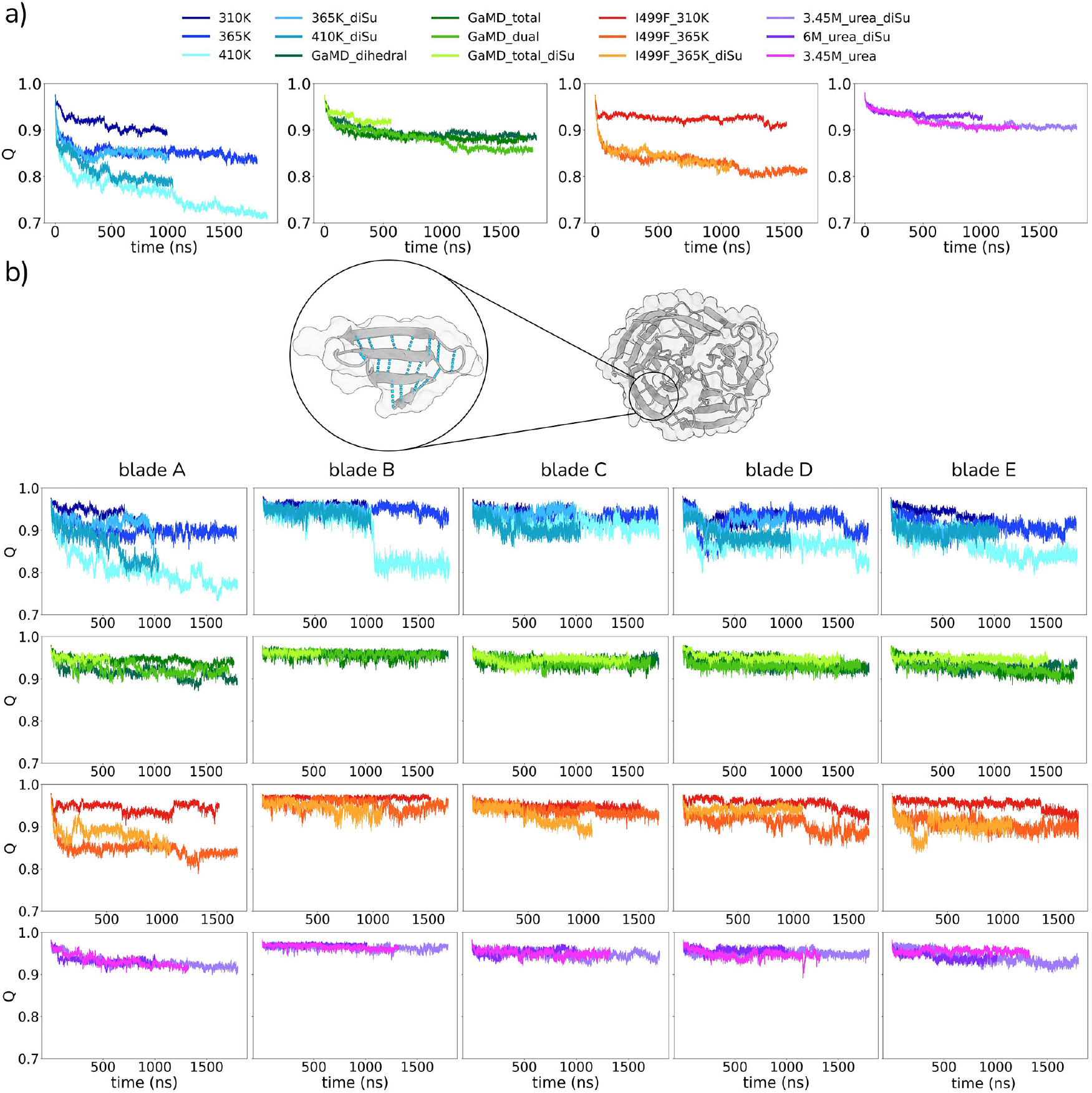
The fraction of native contacts (Q) calculated with soft cutoff was given for a) the entire OLF and b) five blades with reference to the initial OLF structure.

The largest reduction in the fraction of native contacts (Q) was observed in the 410 K simulation, where Q dropped to 0.7, followed by a decrease to 0.8 in both WT and mutant OLFs at 365 K (Fig. 2a). We also observed a gradual decline in Q with increasing temperature (Fig. 2a). At 310 K, a subtle decrease in Q to 0.9 was reported for the WT OLF. This subtle change was similarly the case in the I499F mutant at 310 K, indicating minimal impact of this severe POAG variant [10] on native contacts over 1 *µ*s. Parallel to these subtle changes, a slight expansion of the surface area of WT and I499F OLFs was observed (Fig. S4a). Explicitly, the surface area of the WT enlarged from 118 nm^2^ to 126 nm^2^. The mutant OLF, which displayed a smaller surface area (∼ 120 nm^2^) than WT, also expanded to ∼128 nm^2^ at 310 K.

GaMD simulations decreased Q much less dramatically than did 410 K simulation condition. Particularly, dual-boost potential (targeting both dihedral and total energies) produced the most significant change (Q=0.87), while single-boost potentials (dihedral-only or total energy-only) yielded marginally higher Q values. The disulfide bond partially recovered native OLF contacts in GaMD simulations and WT OLF at 410 K, but had no effect at 365 K for either WT or mutant. In line with its reduced mobility (Figs. S2-S3) and stable SASA profile (Fig. S4), urea solvation did not alter native contacts regardless of concentration or disulfide bond state.

We also traced the native contacts for each blade (Fig. 2b). Consistent with the observations for the entire OLF, 410 K simulations resulted in the largest change in the native contacts within the blades (Fig. 2b). The I499F mutation alone did not induce a notable loss of native contacts within the blades, exhibiting a level of contact reduction (∼ 10%) comparable to that observed in WT OLF at 310 K. However, the I499F mutant at 365 K showed a decline in the native contacts of blades with blade A being the most affected one (Fig. 2b). In urea-solvated systems, all blades preserved over 90% of their native contacts. GaMD simulations, following the urea simulations, showed restoration of native contacts to a level of 90% in all blades for 1 *µ*s. For all simulations, the terminal blades, A and E were the most affected blades.

The *β*-propeller in OLF fold lacks a typical hydrophobic core, instead its structure is stabilized by hydrophobic interactions between the blades (Fig. 1). To monitor how inner blade interfaces were affected, we computed the fraction of native contacts at the blade interfaces (Fig. S5). In all tested MD conditions, the interfaces between blades A-E and A-B were disrupted the most, exhibiting a substantial loss of inter-blade contacts (Fig. S5). The blade interface A-E was almost completely lost in WT and mutant simulations at elevated temperatures (365 or 410 K). The blade A-B interface were the second most affected interface, that was also disrupted in both WT and mutant systems at elevated temperatures. All three GaMD simulations, boosting dihedral, total, or both dihedral and total potential energies, resulted in a reduction of inter-blade contacts between A and E (Fig. S5). In addition to the A-E interface, the interface formed by blades A and B also exhibited a decrease in native contacts in all GaMD simulations. One notable point, the A-E blade interface was intact in the GaMD simulation that applied a boost potential to the total potential energy and also maintained the C245–C433 disulfide bond. Urea solvation affected the native contacts at the A-B interface the most, while other interfaces remained largely intact, maintaining approximately 90% of their native contacts (Fig. S5). The interfaces between the blades B-C, C-D, and D-E generally preserved their native contacts, except for the high-temperature simulation conditions.

These results highlight that the interface formed by the blade A, explicitly A-E and A-B, underwent the most significant contact losses, whereas the B-C interface exhibited the highest stability, followed by C-D and D-E. The substantial loss of the contacts at the A-E interface, which contains the molecular clasp that is essential for maintaining the OLF shape (Fig.1), indicates unfolding of the clasp. Altogether, we reported that the tested MD regimes led to a predominant loss of native contacts at the terminal blade A, either its contacts within itself or with other blades, suggesting its sensitivity to MD induced perturbations and possibly a role in aggregation pathway of OLF.

### Monitoring the Structural Integrity of C245-C433 and Metal Coordination

Whilst exploring possible conformations relevant to aggregation-prone OLF intermediates, we reported a contribution of the disulfide bond formed between C245-C433 to the structural dynamics of OLF (Fig. S2). To characterize the dynamics of these cysteines in the absence of an explicit disulfide linkage, the distance between C245 and C433 was monitored for the simulations where this bond was not defined (Fig. S7). In simulations of I499F mutant, a stable proximity between C245 and C433 was maintained, with the C245-C433 distance reminiscent of a formed disulfide bond, even at 365 K. However, other systems exhibited larger cysteine-cysteine separations in the absence of the bond, frequently exceeding 6 Å (Fig. S7a).

The structural contribution of the C245–C433 disulfide bond to the OLF domain was further investigated by analyzing 1D RMSD profiles of the WT OLF at 410 K, the condition exhibiting the most pronounced structural changes (Fig. 3). In the absence of the disulfide bond, the WT system sampled three distinct RMSD clusters, with gradually increasing RMSD values after the midpoint of the simulation. However, the same condition (410 K) in the presence of the disulfide contact showed a more stable RMSD profile. Evident from the corresponding snapshots from 410 K simulation of WT, the presence of disulfide contact contributed to retain the globular shape of OLF, while in its absence temperature-induced unfolding is more evident with an opening in the structure. Although this structural opening was not observed in the disulfide-stabilized system, a noticeable distortion of the WT OLF shape at 410 K was still present, impacting all inter-blade contacts (Fig. S4). As we also checked the WT OLF system at 365 K, a larger separation of the C245-C433 distance was reported (Fig. S7a) in the WT than mutant, suggesting a contribution of I499F to the integrity of the same region stabilized by the C245-C433 bond. These findings highlight the critical role of the C245–C433 disulfide bond in shaping the OLF structure through restraining its conformations at elevated temperatures.

**Fig. 3.**
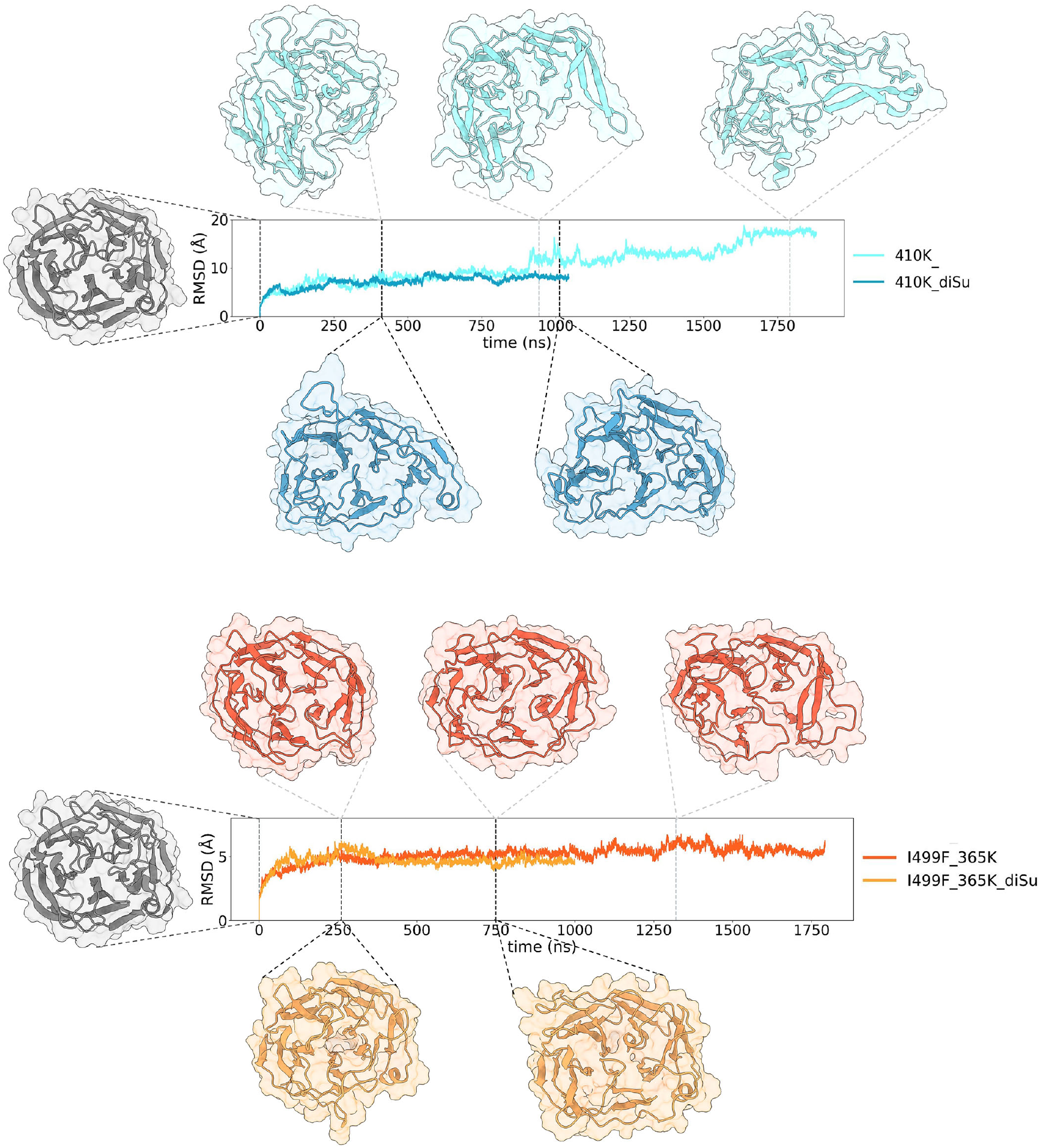
1D RMSD calculations of the WT 410 K (top) and I499F at 365 K (bottom) simulations were given along with representative snapshots at the indicated time points.

We measured the distances of two metal ions, Ca^2+^ and Na^+^, to the C*α* atom of D478 (Fig. S7b-c). Both metals were retained in the central hydrophilic cavity in urea-solvated systems while they were diffused into bulk for all other simulations. We also measured the compactness of hydrophilic cavity by *R*_*G*_ of the residues lining up the dimetallic center (Fig. S8). In line with the distance measurements (Fig. S7b), the radius of the central cavity was stable in the urea-solvated OLF systems. GaMD simulations, WT and I499F OLF at 365 K led to subtle changes in the radius of hydrophilic core, while 410 K simulations of WT OLF greatly expanded the radius, indicating a disruption of the the hydrophilic core.

### Structural Changes Induced by Urea-solvation

To assess the spatial distribution of urea molecules around the protein, we calculated the radial distribution function (RDF) between the centers of mass of urea and the protein (Fig. S9). For all systems, the RDF profiles showed a characteristic exclusion zone below 2 Å and distinct peaks corresponding to the first and second solvation shells between 2 Å and 6 Å, before gradually approaching bulk behavior at larger distances. Notably, the g(r) values remained below 1, indicating hydration of the OLF domain.

Two key differences between the urea solvated systems were apparent. First, the absence of the C245-C433 disulfide bond led to a greater accumulation of urea around the protein surface compared to disulfide bond containing systems. Second, in contrast, increasing the urea concentration did not change the RDF profile significantly. This similarity indicates that higher urea concentration did not alter solvent organization around the protein, consistent with the counterintuitive observation that it did not induce any notable changes in the native contacts of OLF (Fig. 2).

### Predicting Aggregation Propensity of OLF Structures from Different MD Settings

To identify representative conformations from our simulations, we constructed a free energy landscape (FEL) by projecting the dynamics trajectory onto the first two principal components (PC1 and PC2). FEL analysis, which estimates the free energies of sampled states yielded 30 low energy OLF structures from 15 MD simulations with different settings (Fig. S10). Two structure-based aggregation propensity predictors, CamSol [55] and AggreScan3D [50– 52], were used to analyze these representative MD structures and the initial PDB structure (8frr) (Fig. S11). CamSol predicts solubility and identifies low-solubility regions as aggregation-prone, while AggreScan3D identifies aggregation propensity of each amino acid using an experimental aggregation propensity scale. Though its scores were low, the PDB structure of WT OLF (8frr), readily showed some potential APRs in the N-terminus and blades A and D (Fig. S11). We also constructed the electrostatic potential distributions of the selected structures along with the PDB structure (Fig. S12). Particularly, the selected snapshots from high temperature simulations of WT OLF show notable difference from the PDB structure by means of electrostatic surface potentials.

To identify how aggregation propensity changed during our simulations, we calculated the score difference between the selected representative MD structures and the 8frr structure (Fig. 4). For this we selected on set of representative MD structures for each simulation. Positive peaks on *y*-axes of plots in Fig. 4 show regions that became more aggregation-prone in the MD snapshots while negative values indicated regions where the aggregation propensity decreased or remained the same.

**Fig. 4.**
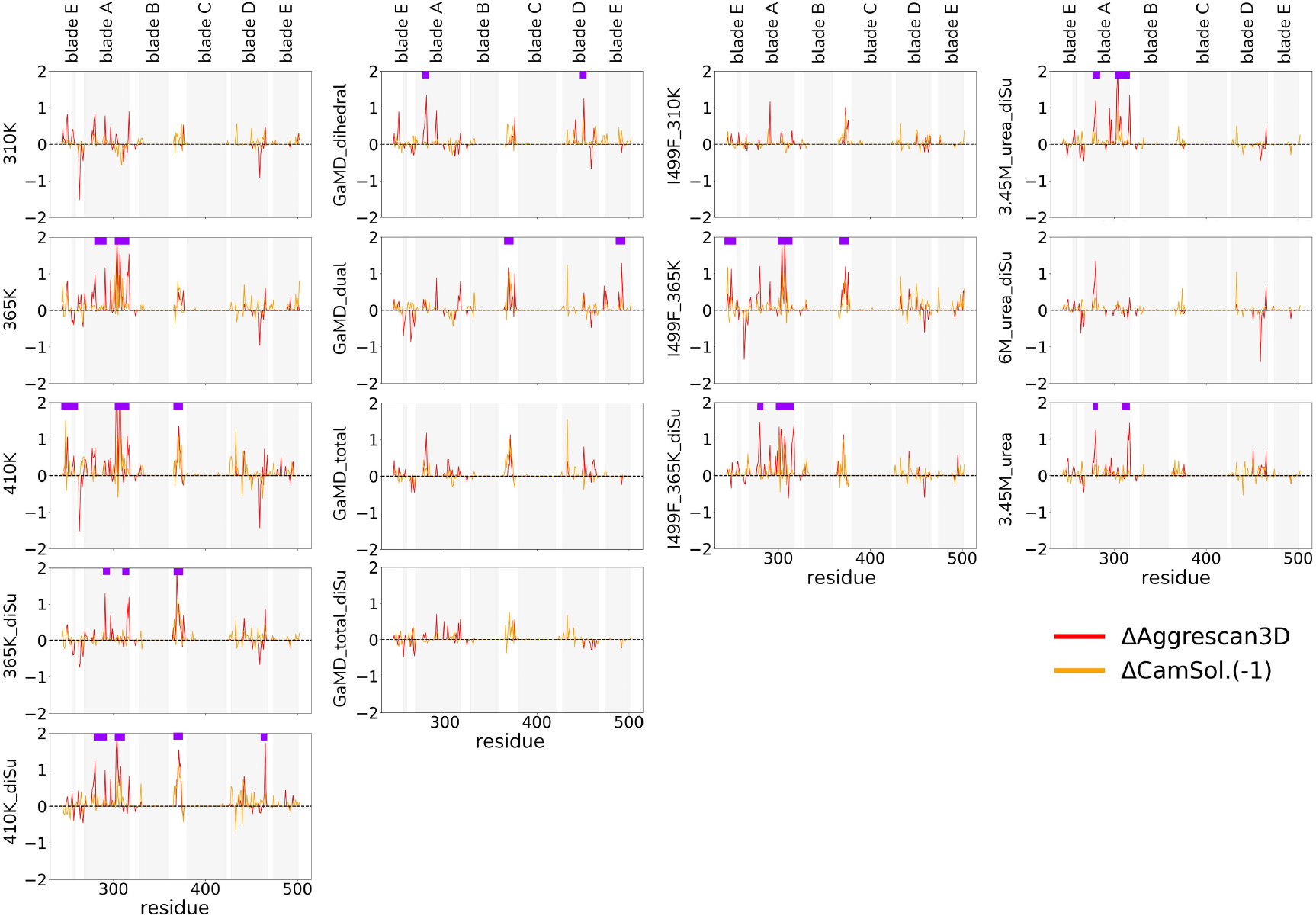
Changes in aggregation propensity referencing the 8frr structure calculated by AggreScan3D and CamSol were given.

For the representative snapshots from the WT and I499F simulations at 310 K, the aggregation propensity remained largely unchanged, with no regions showing peaks greater than 1 unit compared to the static structure (Fig. 4). On the other hand, representative structures from all other simulations displayed at least one region with a score difference exceeding 1 unit for both methods. Notably, blade A consistently showed a marked increase in scores in nearly all MD conditions. Additionally, the N-terminal region of the OLF domain corresponding the connecting region between blades A and E also showed larger score deviations, particularly in the 410 K simulation of the WT and the 365 K simulations of the I499F mutant. We observed a gradual increase in the aggregation propensity of blade A and the long loop between blades B and C, with increasing temperature for both WT and I499F mutant simulations. The urea simulation also showed increased aggregation tendency specifically in the blade A region. In the GaMD simulations, we observed aggregation potential, particularly for the long loop and blades A, D and E. While none of the representative structures showed an increase in aggregation propensity scores for the blades B and C.

We visualized the regions showing a large positive differences in aggregation propensity relative to the 8frr structure (Fig. 5). Three regions in blade A, which particularly showed higher aggregation propensity for high temperature simulations of both WT and I499F mutant, are ^275^KPTYPYTQE^284, 288^IDTVGTDVR^296^ and ^301^YDLISQFMQGPSKVHIL^317^. Among these, ^288^IDTVGTDVR^296^ was also appeared in the GaMD and urea solvated simulations (Fig. 5). For all three regions, short patches of hydrophobic amino acids exhibited high aggregation propensity (Fig. 4). Essentially, these include Y280 for the first region; I291 for the second region; Y301, L303, I304, F307, M308, V314, H315, I316, L317 for the third region. Visualization of the corresponding MD snapshots suggested that these hydrophobic amino acids became more accessible in the MD snapshots compared with the static PDB structure. Particularly, the third region is tightly packed and folded into a short helix in the PDB structure while the MD conditions led to structures that lost this helix rendering hydrophobic amino acids within this helix accessible. Another region that also recurrently appeared in distinct MD simulations was located within ^246^GELVWVGEPLTLR^259^ corresponding to the molecular clasp (Fig. 4). This region was appreared in the 410K simulation of WT with L248 and W250 residues showing high scores (Fig. 4). Another region, ^364^GYHGQFPYSWG^374^, was also emerged in the high temperature simulations of both WT and mutant along with GaMD simulation with boost potential applied to both total and dihedral potential energies. Similarly, a short hydrophobic patch in this region (370-374) showed high scores. Two other regions (^460^SKTLTIPF^467^ and ^488^AWDNLNM^494^) were also appeared in the blades D and E (Fig. 5). The region between 460-467 was also recovered from the 410K simulation of WT OLF while the region between 488-494 in blade E appeared in the one of the GaMD simulations. For both regions, two hydrophobic amino acids, I465 and L492, displayed high scores (Fig. 4). Overall, structural visualization of the representative snapshots from MD systems suggested possible aggregation prone region in OLF structure which are rich in hydrophobic and/or aromatic amino acids.

**Fig. 5.**
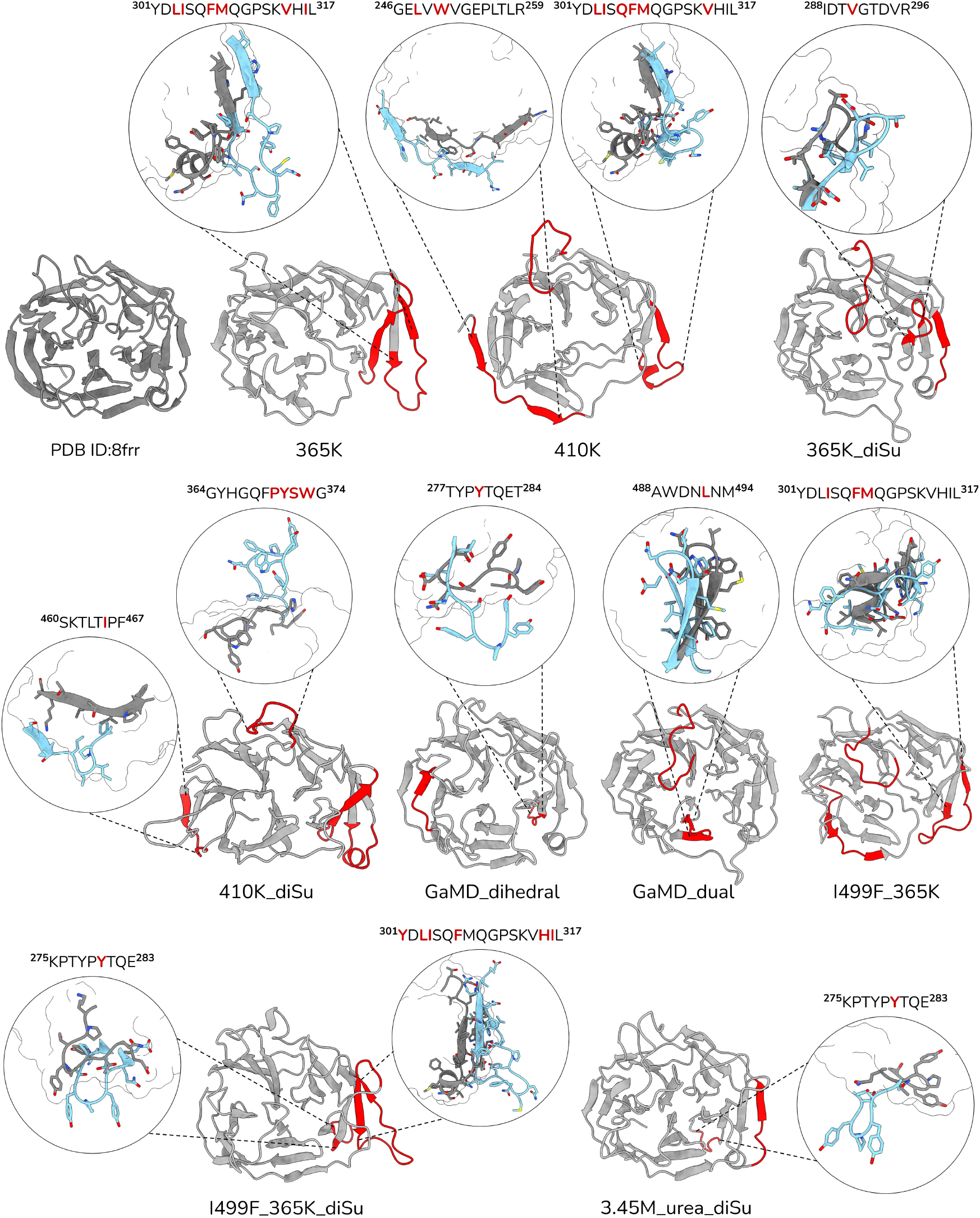
Aggregation-prone regions and changes in side chain orientations were depicted on the OLF conformations from various settings. The sequence of APR peptides was also given.

## Discussion

Slow dynamics of OLF have posed a challenge for classical MD simulations to capture aggregation-prone intermediates of OLF related to POAG pathology [10, 25, 57]. Our study overcomes this barrier by employing a multi-faceted simulation strategy. By integrating unbiased enhanced sampling (GaMD), high-temperature thermal stress, chemical denaturation with urea, and the study of the pathogenic I499F mutant, we populated and characterized potential intermediate states.

Previous experimental studies have established that under mildly destabilizing conditions, the OLF domain of myocilin unfolds and aggregates, a process initiated by the exposure of its buried internal regions [9, 10, 21]. To investigate the structural basis of this chemical-induced aggregation, we simulated the OLF domain in a urea solution at 310 K, mimicking established experimental conditions [10]. While the overall fold of OLF remained intact, our simulations revealed a specific opening of the interface between blades A and B, resulting in the loss of ∼ 50% of their contacts (Fig. S4). This subtle structural change provides a mechanism for the experimental observations. The separation of blades A and B exposes a buried region, ^275^KPTYPYTQE^284^, which consequently shows an increase in aggregation propensity in the representative MD structures (Figs. 4 and 5). Specifically, Y280, which is initially buried (SASA =19 Å^2^, relative SASA=0.07 based on [58]), becomes fully exposed. In turn, aggregation propensity predictors identify the newly accessible Y280 as an aggregation hotspot. Thus, we appraise that the destabilization of the A-B blade interface which lead to the exposure of an aggregation-prone region, proposing a structural explanation for how urea-led unfolding initiates the aggregation cascade of OLF.

Crucially, this mechanism is not unique to urea solvation. We observed the same destabilization of the blade A-B interface and subsequent exposure of the 275-284 region in simulations of the disease-associated I499F mutant and in accelerated MD simulations (Figs. 5 and S4). In all cases, Y280 consistently become solvent exposed and emerged as one of the most aggregation-prone residues. The repeated identification of this region harboring Y280 in multiple simulations pinpoints the interface between blades A and B as a critical region for aggregation of WT OLF.

In addition to this APR at the blade A-B interface, our simulations consistently identified a second APR on the exterior of blade A contacting blade E (^301^YDLISQFMQGPSKVHIL^317^). This external APR at the interface of blades A and E showed high amyloidogenic propensity in multiple simulations, including high temperature and urea solvation. The POAG-associated E323K mutant, which trails this APR, is found at the blade A-B junction [59]. Notably, the external region encompassing 301-307 and the previously discussed internal region (275-284) together flank the P1 peptide (326-337), an OLF segment experimentally shown to be highly amyloidogenic [60]. While the P1 peptide itself did not exhibit high aggregation propensity in our MD structures, its close position to two potential APRs is noteworthy. We propose a mechanism whereby the destabilization of these two flanking regions (275-284 and 301-317), starting from the exposure of the aromatic YPY motif in the internal region, initiates the conformational changes necessary to unmask the P1 peptide. Altogether, this model provides a structural pathway that connects our computational findings to experimental results, suggesting that these two regions could be upstream triggers for P1-mediated OLF aggregation.

Our simulations also revealed that the short side helix inside the region 301-317 is a point of structural lability. Under thermal denaturation stress as well as in GaMD simulations, this helix transitions to a disordered state for both WT and I499F mutant (Fig. 5). This transition affects the overall compactness of OLF (Fig. 5) and in turn contributes to the increased aggregation propensity scores for the ^301^YDLISQFMQGPSKVHIL^317^ region (Fig. 4). Hence, integrity of this interface and the short helix is critical for maintaining a stable, non-aggregating conformation. We also note that the observed instability from simulations is consistent within the experimental OLF structures. Essentially, a survey of PDB entries of OLF domain reveals that this short helix is disordered in some structures. In one particular case (PDB id: 6pkf), this surface shows a completely different helical arrangement, adopting a single-turn helix in blade E, instead of blade A, highlighting its inherent flexibility in the experimental structures of OLF as well. Therefore, we identify the A-E interface and the short helix as another aggregation hotspot, where its order-to-disorder transition can lead to exposure of regions contributing to the aggregation cascade.

The C245-C433 disulfide bond is a critical stabilizer of the OLF domain’s five-bladed *β*-propeller structure [11, 24]. By covalently linking the N-terminal loop to the surface of blade D, this bond acts as a crucial structural tether particularly for the molecular clasp (Fig. 1). Our simulations confirm that this linkage is essential for maintaining the OLF toroidal fold, conferring resistance particularly under thermal denaturation stress (Fig. 3). In its absence, the domain experiences significant structural perturbations, leading to the loss of toroidal shape (Fig. 3). This observation also support the structural mechanism for why its disruption, as seen in the pathogenic C433R variant, leads to loss-of-function and a high propensity for aggregation [25, 61]. Importantly, our analysis of the POAG-associated I499F mutant reveals how other interactions can compensate for the loss of this disulfide bond. While the WT OLF shows large structural perturbations at elevated temperatures without the C245-C433 linkage, the I499F mutant remains stable under similar conditions, preserving its native-like state. This is due to enhanced local packing of an internal hydrophobic cluster. Essentially, the substitution of I with a bulkier F at the position 499 stabilizes a hydrophobic cluster formed by L486, I432, Y479, L501 within the blades D and E. This cluster appears to act as an alternative anchor, mimicking the stabilizing role of the disulfide bond. Overall, these observations suggest an interplay between the C245-C433 disulfide bond and hydrophobic interactions in preserving the integrity of the OLF domain, reflecting the significance of the redox state of the C245-C433 in investigating OLF intermediates.

Another aggregation-prone region was emerged in the long loop (364-374) in our high-temperature and enhanced sampling simulations, with the hydrophobic PYSW motif showing particularly high aggregation scores (Fig. 4). This APR contains P370, the site of the critical P370L mutation known to cause severe, early-onset glaucoma [62, 63]. Our analysis therefore further suggests a mechanistic basis for the pathogenicity of P370L, a destabilizing mutation [25, 61], located in a region that is susceptible to aggregation. This insight, combined with our findings for other critical sites such as C433R and E323K variants, exemplifies how our approach can link disease-causing mutations to specific points of structural vulnerability in the protein.

Our simulations revealed a paradoxical effect where the OLF domain remained almost completely intact at high urea concentrations, while showing initial signs of unfolding, particularly for the first APR between 275-284 at a lower urea concentration (Fig. 5). Unequivocally, this not a genuine stabilization at high urea concentration, but an artifact probably arising from timescale limitations of classical MD simulations [64, 65]. A notable similarity was observed between two RDFs at high and low urea concentrations (Fig. S9), implying that overall solvent organization remained nearly identical in both simulations. If the 6M urea concentration were actively inducing denaturation, a significantly different RDF profile would be expected. Instead, our results suggest that OLF is trapped in a caged state at high urea concentration. Furthermore, this unexpected stability of OLF (Fig. 2) could also be a result of urea self-aggregation and/or drastic increase in solvent viscosity at high urea concentration, limiting the direct denaturing interactions of urea with the protein [66]. We overall report that classical MD simulations at 6M urea concentration failed to capture the anticipated unfolding behavior of OLF [10]), underscoring the challenging nature of OLF folding-unfolding dynamics.

## Conclusions

By integrating enhanced sampling with classical molecular dynamics simulations under diverse stress conditions, we navigated the challenging conformational landscape of the myocilin OLF domain, producing a comprehensive structural map of the amyloidogenic regions of OLF. Identification of seven distinct APRs, several of which were previously elusive, are partly validated by the experimental data on the amyloid-forming peptides and the POAG variations. The main insight of this study is, thus, a detailed model where OLF aggregation is not random, but a structured process initiated at specific regions. Our findings can pave the way for future directions. First, the APRs identified here serve as immediate targets for experimental studies, e.g. via site-directed mutagenesis, to confirm their aggregation-driving roles. Second, the structural map documented here provides a blueprint for the design of small molecules or peptides or antibodies aimed at binding/shielding the APRs to halt the amyloid cascade [67]. Ultimately, the multi-faceted MD simulations strategy employed in this study offers a powerful and transferable framework for investigating challenging amyloid systems.

## Acknowledgments

All of the calculations reported in this paper were performed by the computational resources at TUBITAK ULAKBIM, High Performance and Grid Computing Center (TRUBA resources).

## Data Availability Statement

OLF conformations from various settings with aggregation-prone regions are available at https://doi.org/10.5281/zenodo.16712800. OLF MD simulation analysis scripts are available at https://github.com/timucinlab/OLF_apr_search.git.

