## Supplementary File for "Aggregation-Prone Region Mapping in Olfactomedin Domain of Myocilin through Classical and Enhanced Sampling Molecular Dynamics Simulations"

### Supplementary Information

#### Supplementary Tables

**Table S1.** MD System Details

| id | protein | water | urea | ion† | x, y, z (Å) | time (ns) |
|---|---|---|---|---|---|---|
| 310K | 4077 | 27771 | 0 | 57 | 70, 70, 71 | 995 |
| 365K | 4077 | 27786 | 0 | 57 | 70, 70, 70 | 1794 |
| 365K_diSu | 4075 | 27780 | 0 | 57 | 70, 70, 70 | 1000 |
| 410K | 4077 | 27798 | 0 | 57 | 71, 70, 70 | 1881 |
| 410K_diSu | 4075 | 27780 | 0 | 57 | 70, 70, 70 | 1044 |
| GaMD_dihedral | 4077 | 27771 | 0 | 57 | 70, 70, 71 | 1800 |
| GaMD_dual | 4077 | 27771 | 0 | 57 | 70, 70, 71 | 1721 |
| GaMD_total | 4077 | 27771 | 0 | 57 | 70, 70, 71 | 1768 |
| GaMD_total_diSu | 4075 | 27804 | 0 | 57 | 70, 70, 70 | 566 |
| I499F_310K | 4078 | 27795 | 0 | 57 | 70, 70, 70 | 1519 |
| I499F_365K | 4078 | 27792 | 0 | 57 | 70, 70, 71 | 1681 |
| I499F_365K_diSu | 4076 | 27786 | 0 | 57 | 70, 70, 70 | 1080 |
| 3.45M_urea_diSu | 4073 | 33000 | 9600 | 82 | 80, 80, 80 | 1825 |
| 6M_urea_diSu | 4073 | 30000 | 19200 | 82 | 80, 80, 80 | 1013 |
| 3.45M_urea | 4075 | 33000 | 9600 | 82 | 80, 80, 80 | 1323 |

†All systems were neutralized to a final NaCl concentration of $0.15M$.

#### Supplementary Figures

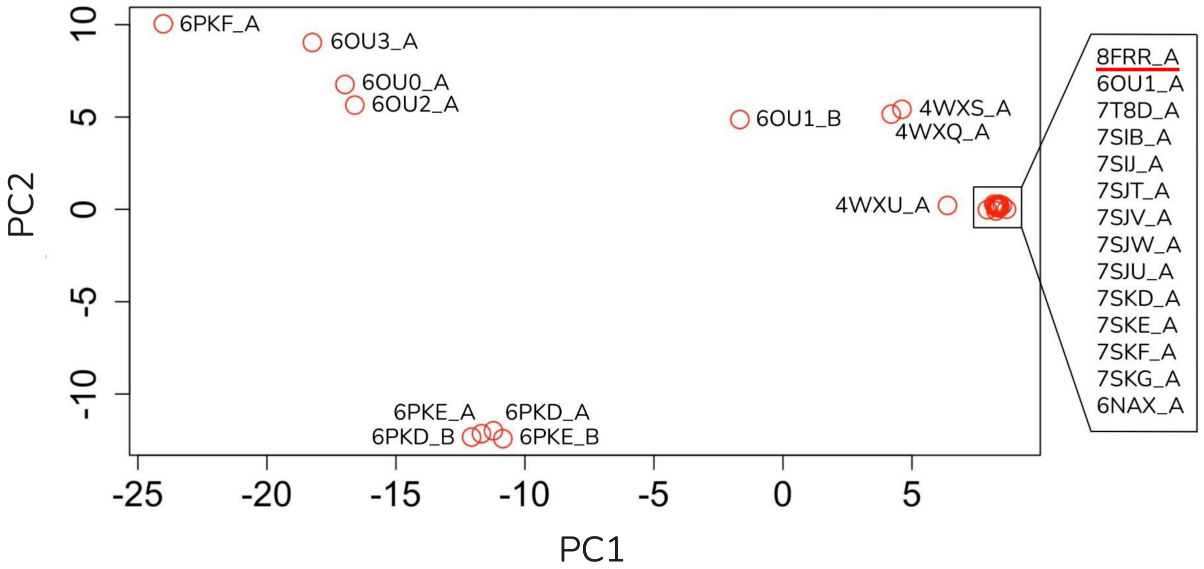


**Fig S1.** PC1–PC2 score plot from principal component analysis (PCA) based on the C$\alpha$ coordinates of 26 experimental structures of the human myocilin OLF domain available in the PDB as of January 2025. Among these, the structure with PDB ID 8frr was selected for this study.

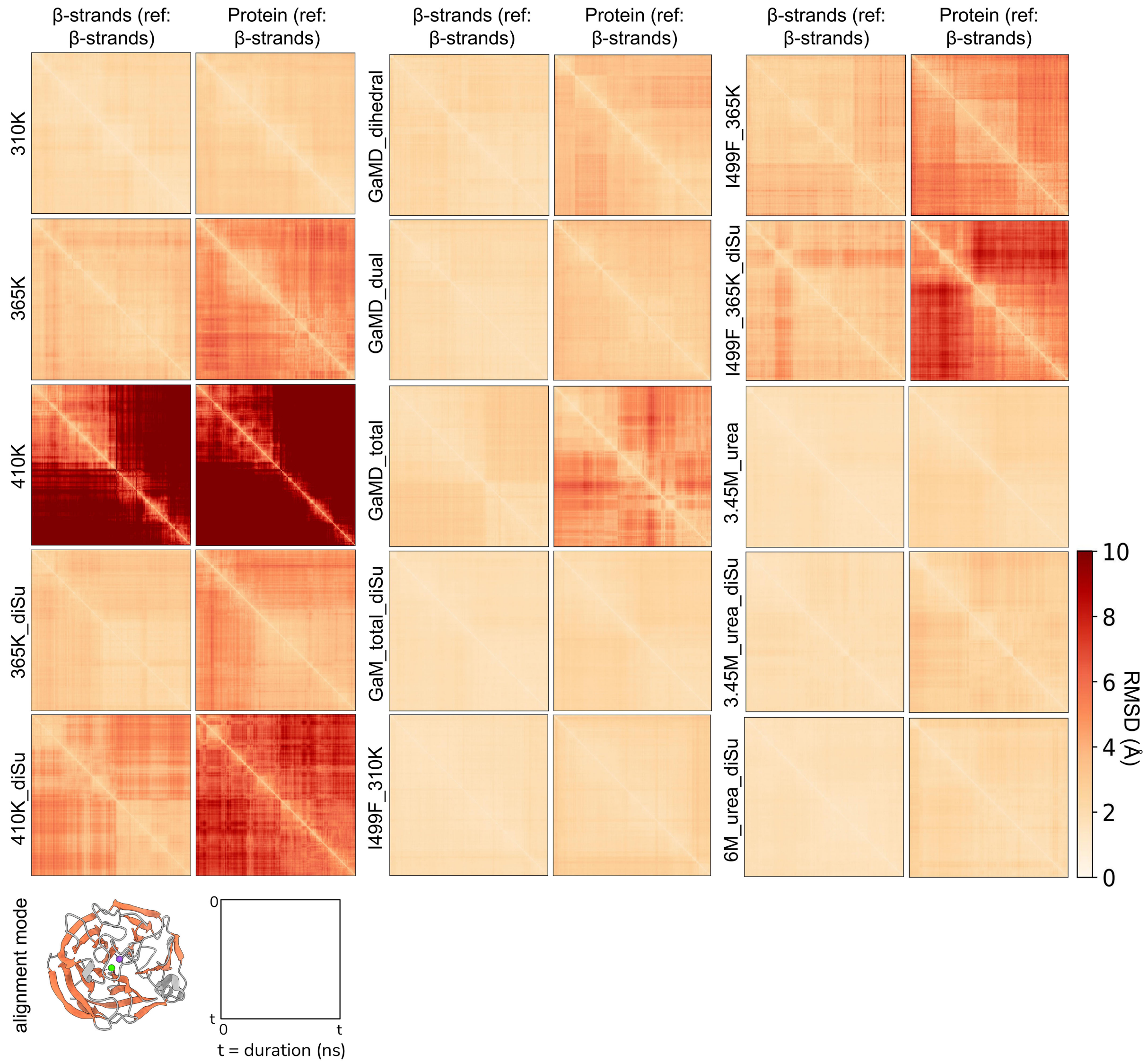


**Fig S2.** Pairwise RMSD analysis of 15 MD simulations. For each system, left panel shows Cα RMSDs of the blades, and right panel shows Cα RMSDs of the entire OLF. For both, the Cα trace of the blades were used as the reference.

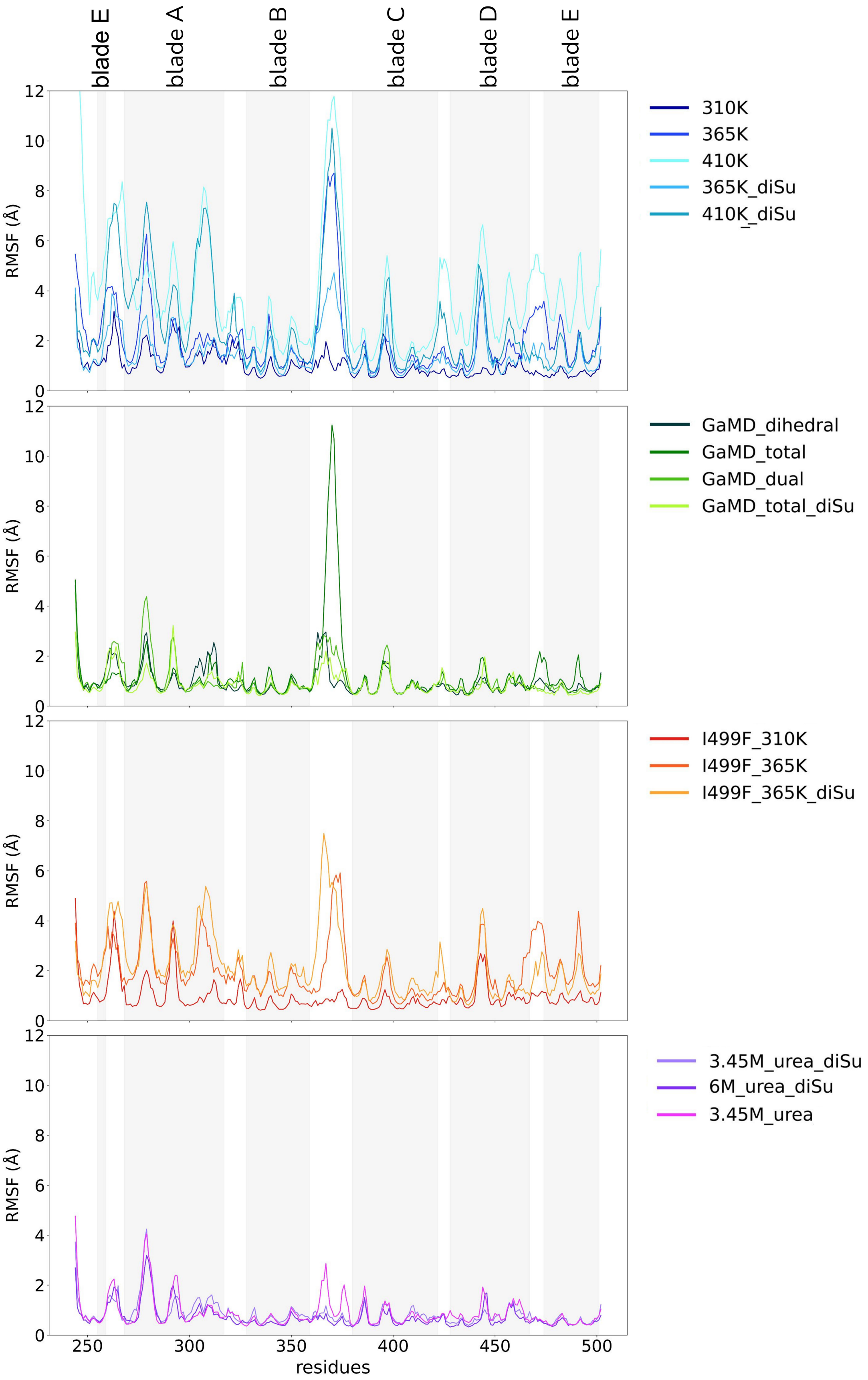


**Fig S3.** C$\alpha$ fluctuations for (a) WT OLF simulations at varying temperatures, (b) WT in GaMD simulations with different boost potentials, (c) I499F mutant at varying temperatures, and (d) WT in urea solutions.

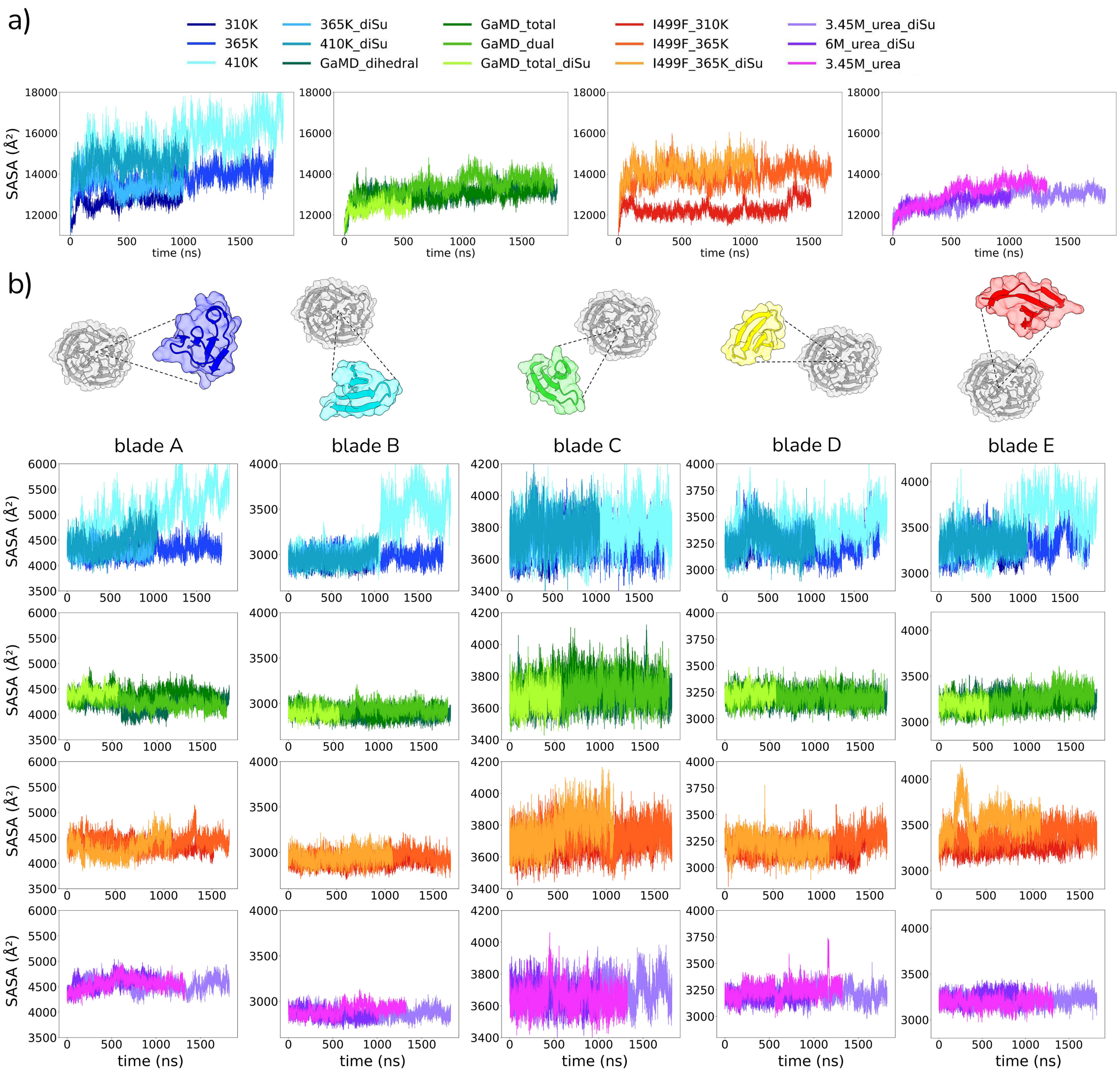


**Fig S4.** SASA measurements for (a) entire OLF and (b) blade structures.

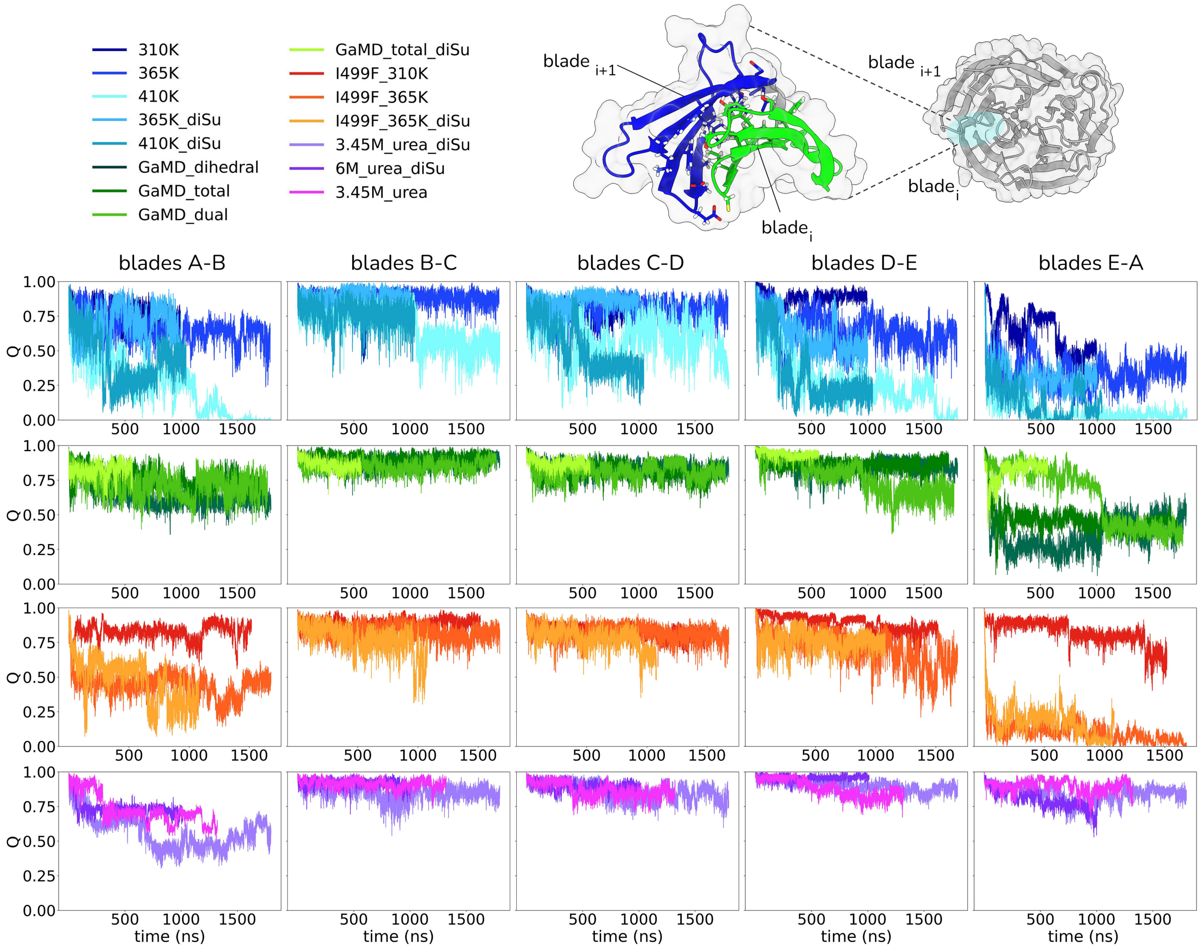


**Fig S5.** Fraction of native contacts (Q) for the inter-blade contacts.

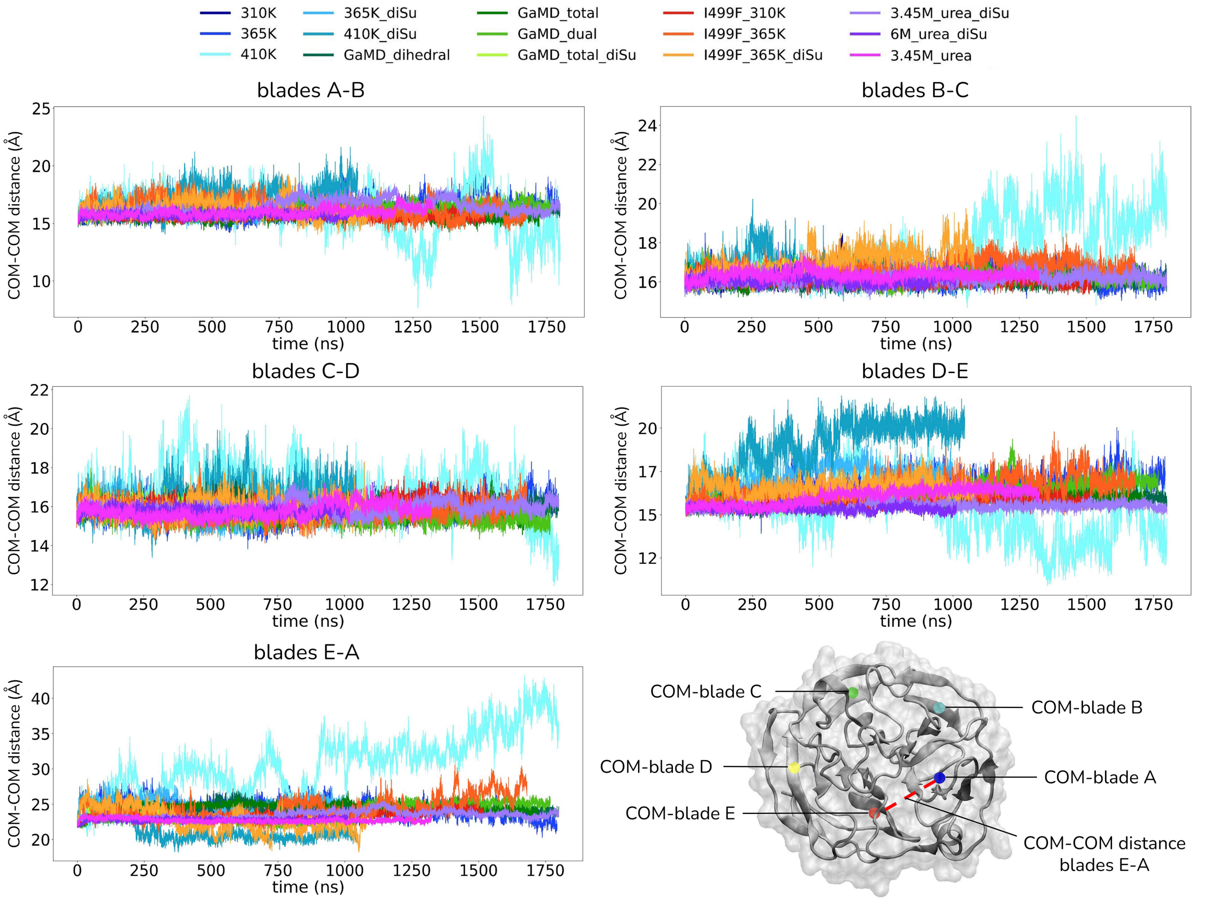


**Fig S6.** Distances between COM-COM of two neighboring blades.

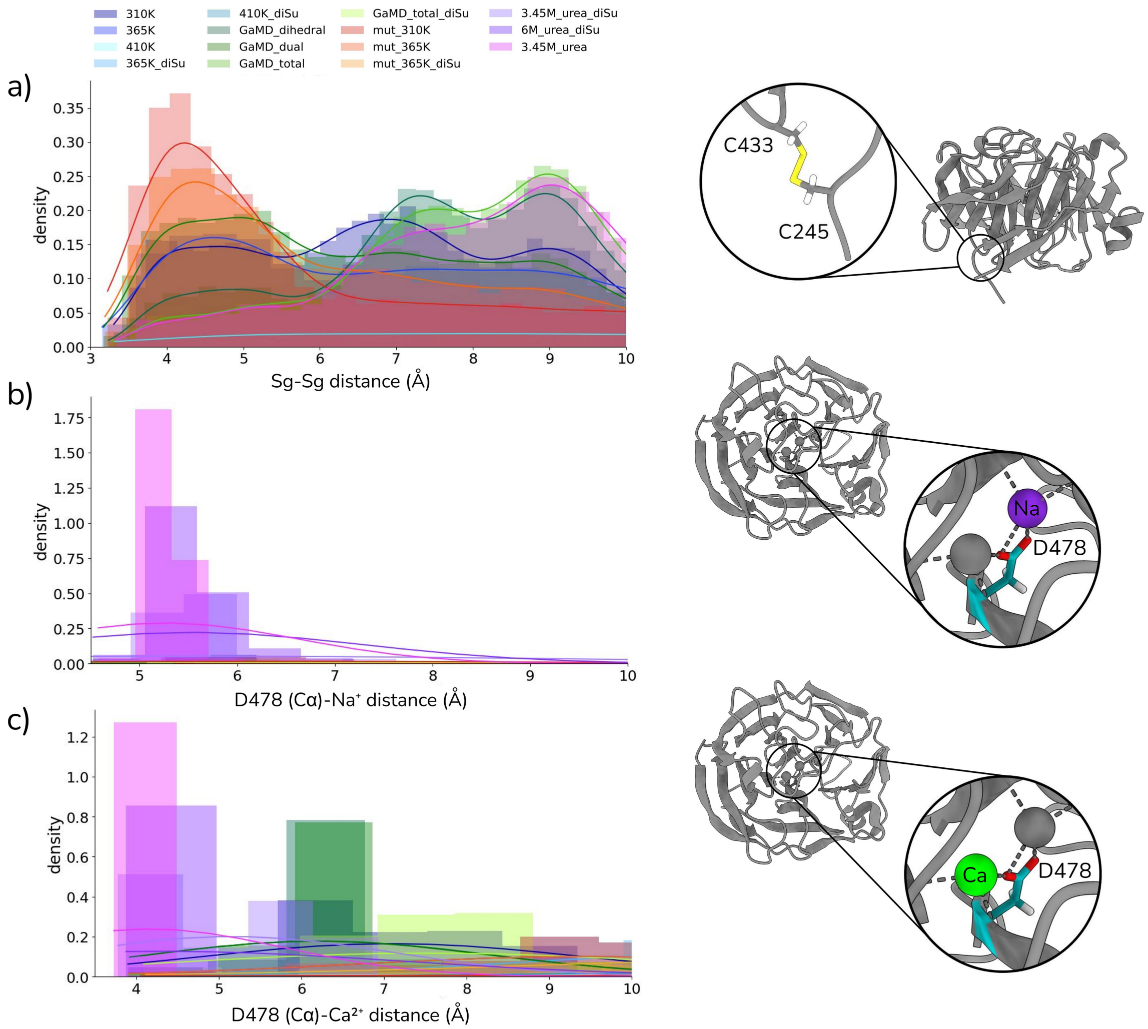


**Fig S7.** (a) Distance between C245-C433 for the systems without the disulfide bond, (b) the distance between sodium and D478 (C$\alpha$) and (c) calcium and D478 (C$\alpha$).

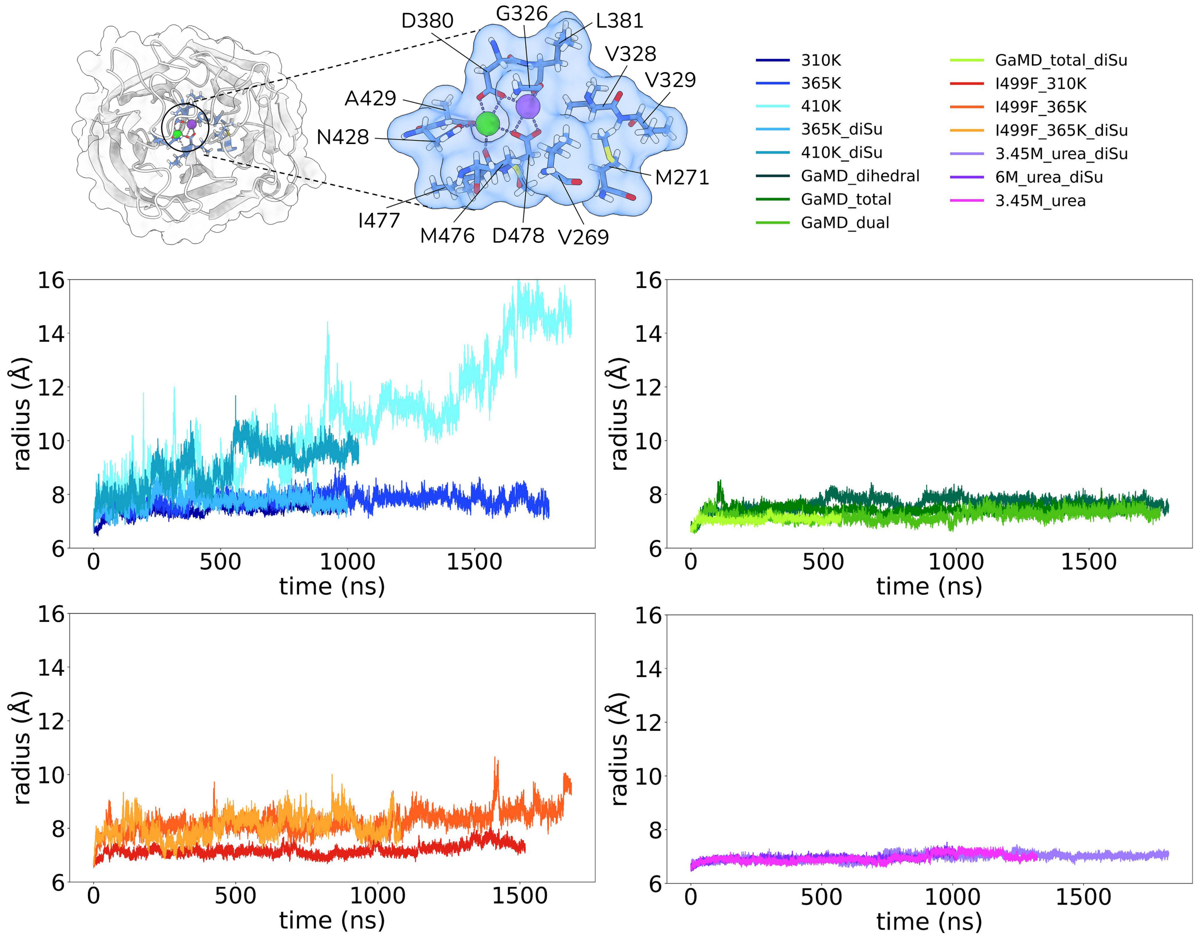


**Fig S8.** Measurement of the radius of gyration ($R_G$) of the residues forming the dimetallic center (V269, M271, G326, V328, V329, D380, L381, N428, A429, M476, I477, D478).

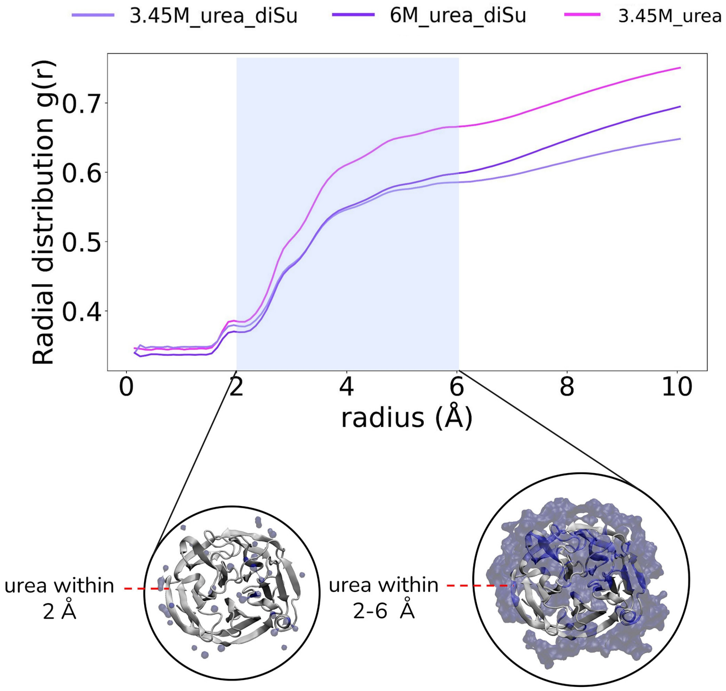


**Fig S9.** COM-COM radial distribution function (RDF) for protein and urea. Urea molecules around the protein surface at 2 Å and 2-6 Å were illustrated for the system at 6 M urea concentration.

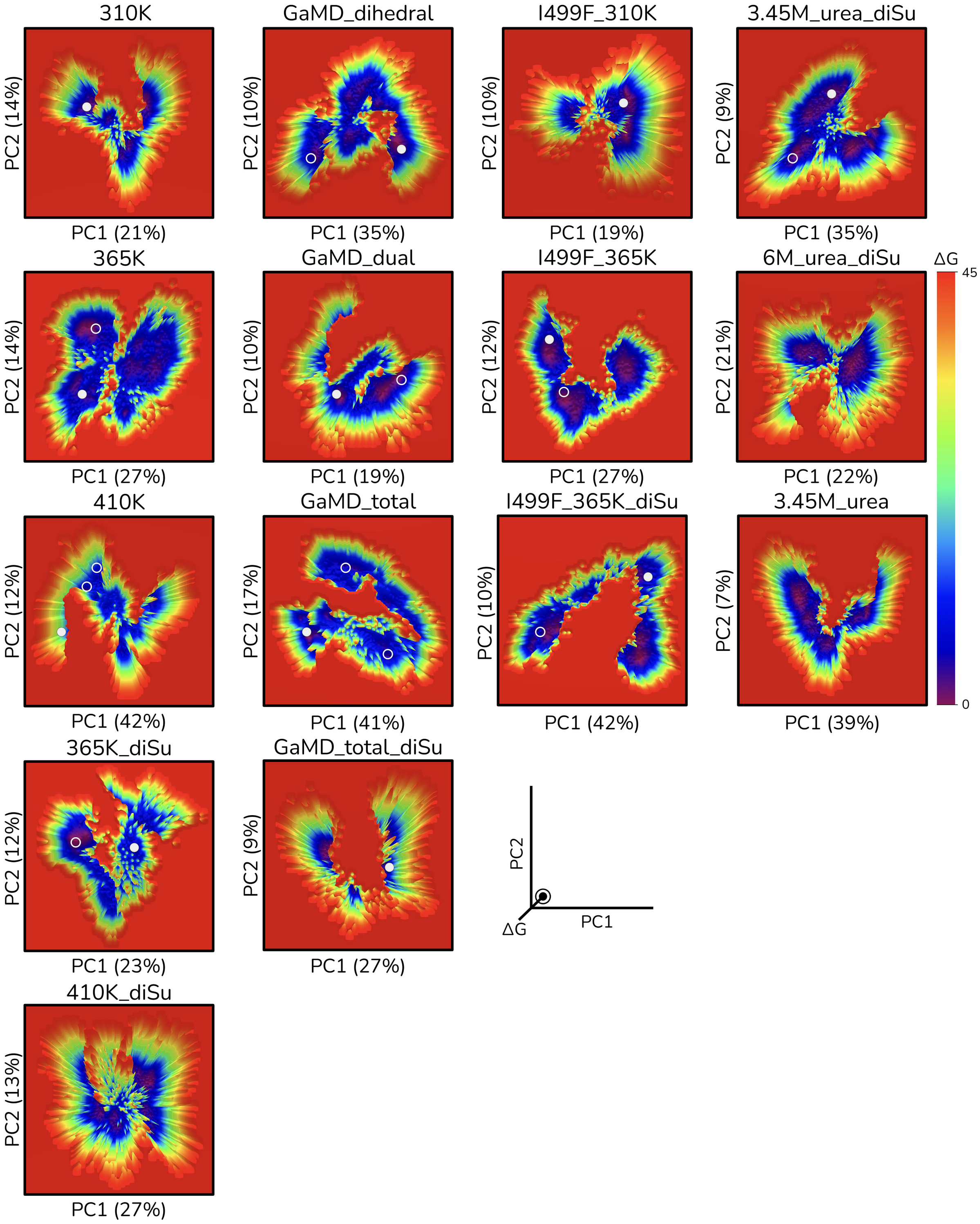


**Fig S10.** Free Energy Landscape (FEL) was projected onto the first two principal components (PC1 and PC2). Each point, representing a sampled structure, colored according to its free energy with purple shades indicating lower energy basins. Selected conformations were marked by circles.

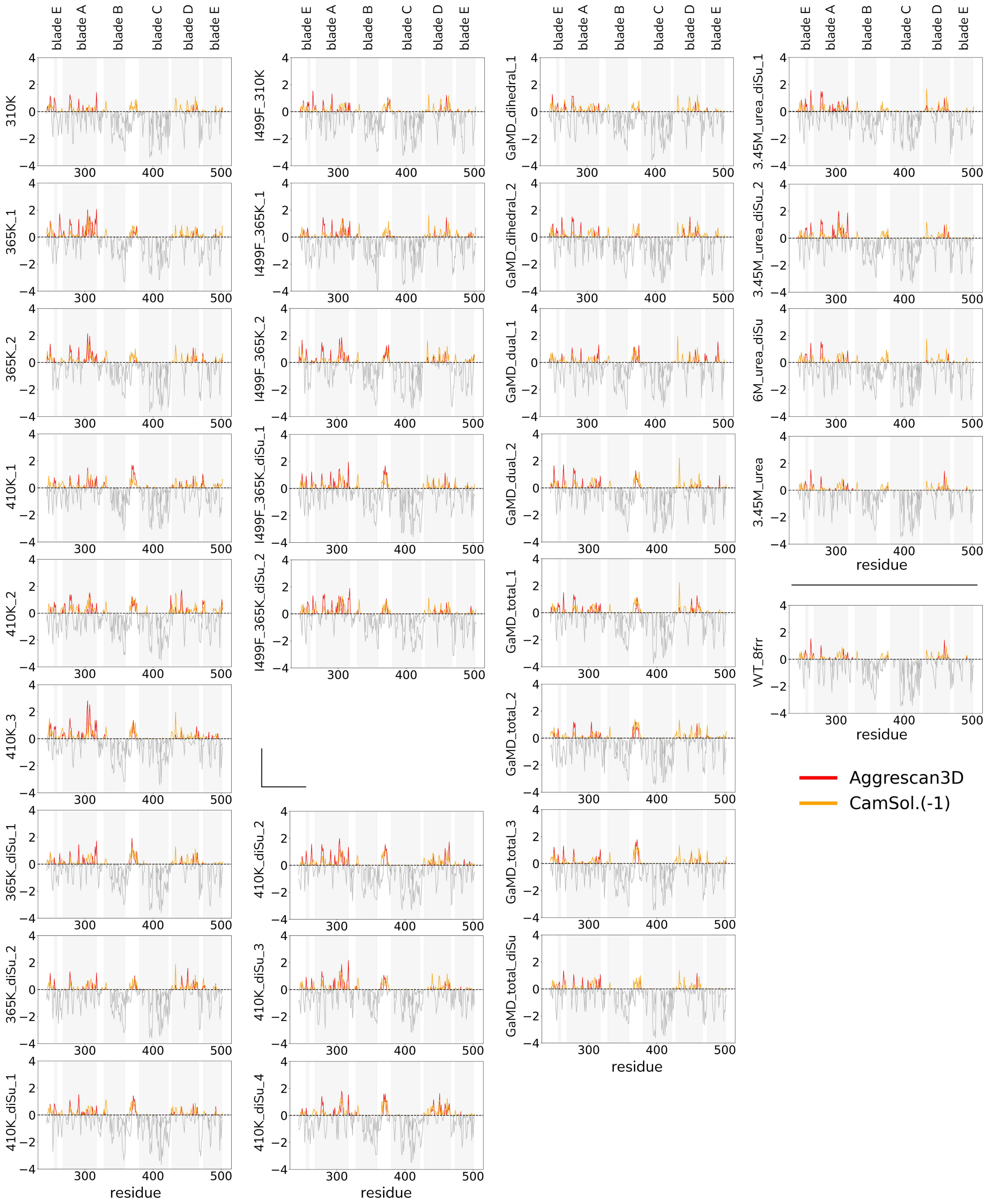


**Fig S11.** Aggregation propensity of the selected OLF structures was predicted using AggreScan3D (red) and CamSol (orange). For clarity, residues with scores below zero were shown in gray. CamSol scores were inverted so that positive values indicate aggregation-prone or insoluble regions.

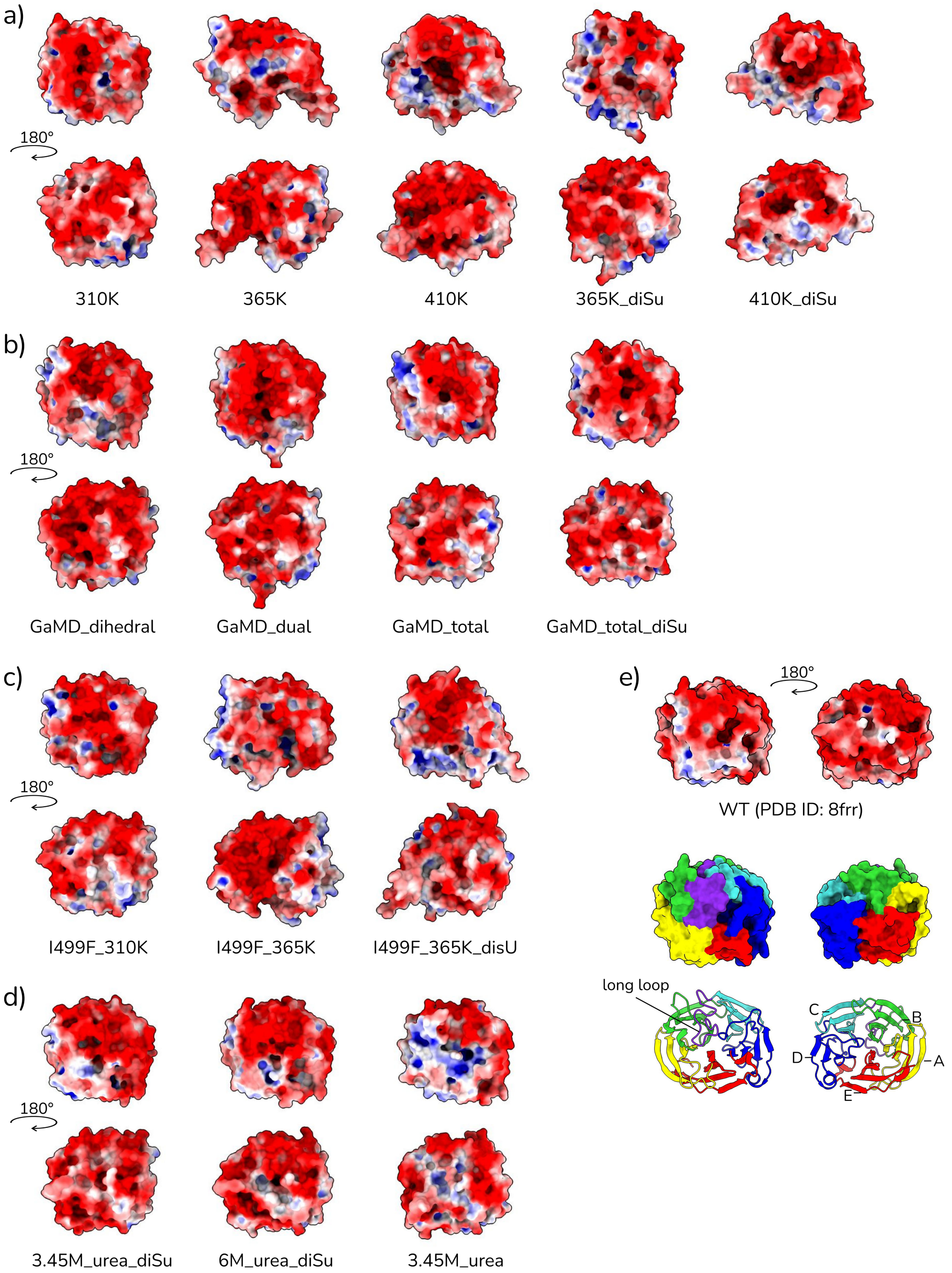


**Fig S12.** Electrostatic potential distributions, calculated using the Adaptive Poisson–Boltzmann Solver (APBS), were mapped onto the molecular surfaces of intermediate conformations selected from (a) WT OLF at various temperatures, (b) WT OLF from GaMD simulations, (c) the I499F mutant at different temperatures, and (d) WT OLF in urea solution. For reference, the surface charge distribution of the initial PDB structure is shown in (e).
